# MVComp toolbox: MultiVariate Comparisons of brain MRI features accounting for common information across metrics

**DOI:** 10.1101/2024.02.27.582381

**Authors:** Stefanie A Tremblay, Zaki Alasmar, Amir Pirhadi, Felix Carbonell, Yasser Iturria-Medina, Claudine J Gauthier, Christopher J Steele

## Abstract

Multivariate approaches have recently gained in popularity to address the physiological unspecificity of neuroimaging metrics and to better characterize the complexity of biological processes underlying behavior. However, commonly used approaches are biased by the intrinsic associations between variables, or they are computationally expensive and may be more complicated to implement than standard univariate approaches. Here, we propose using the Mahalanobis distance (D2), an individual-level measure of deviation relative to a reference distribution that accounts for covariance between metrics. To facilitate its use, we introduce an open-source python-based tool for computing D2 relative to a reference group or within a single individual: the MultiVariate Comparison (MVComp) toolbox. The toolbox allows different levels of analysis (i.e., group-or subject-level), resolutions (e.g., voxel-wise, ROI-wise) and dimensions considered (e.g., combining MRI metrics or WM tracts). Several example cases are presented to showcase the wide range of possible applications of MVComp and to demonstrate the functionality of the toolbox. The D2 framework was applied to the assessment of white matter (WM) microstructure at 1) the group-level, where D2 can be computed between a subject and a reference group to yield an individualized measure of deviation. We observed that clustering applied to D2 in the corpus callosum yields parcellations that highly resemble known topography based on neuroanatomy, suggesting that D2 provides an integrative index that meaningfully reflects the underlying microstructure. 2) At the subject level, D2 was computed between voxels to obtain a measure of (dis)similarity. The loadings of each MRI metric (i.e., its relative contribution to D2) were then extracted in voxels of interest to showcase a useful option of the MVComp toolbox. These relative contributions can provide important insights into the physiological underpinnings of differences observed. Integrative multivariate models are crucial to expand our understanding of the complex brain-behavior relationships and the multiple factors underlying disease development and progression. Our toolbox facilitates the implementation of a useful multivariate method, making it more widely accessible.

## 1. Introduction

In the past decade, there has been exponential growth in the number of modeling approaches that link white matter (WM) microstructural properties and the MR signal (Novikov et al., 2018). Since none of the existing models (e.g., diffusion tensor, neurite orientation dispersion and density imaging (NODDI), etc.) is a perfect representation of the underlying microstructure, choosing a model and contrast for analyses can be challenging, potentially leading to errors in biological interpretation (Novikov et al., 2018). Multi-modal imaging, and multivariate frameworks that combine several parameters derived from different models and modalities, have been suggested as a promising avenue to harness the complementarity of different neuroimaging-derived metrics (Tardif et al., 2016; Uddin et al., 2019).

Multivariate frameworks have the potential to counteract issues arising from the physiologically unspecific nature of commonly used neuroimaging metrics and to capture the complexity and heterogeneity of biological properties (Dean et al., 2017; Guberman et al., 2022; Seidlitz et al., 2018; Tardif et al., 2016; Taylor et al., 2020).Multiple mechanisms give rise to brain structure (e.g., myelination, cell proliferation), support neuroplastic change (e.g., Azzarito et al., 2023; Taubert et al., 2012) and behavioral performance (e.g., Seidlitz et al., 2018; Thiebaut de Schotten & Forkel, 2022), and are involved in neurological disorders (e.g., Iturria-Medina et al., 2017). Interpreting the results of univariate statistical analyses is thus challenging within this context. In addition to capturing a more nuanced picture of the expected mechanisms, multivariate statistical frameworks can offer greater statistical power than multiple univariate analyses as they reduce the amount of multiple comparisons correction required (Avants et al., 2008; Naylor et al., 2014; Owen et al., 2021). Lastly, and perhaps most importantly, multivariate frameworks can be leveraged to move away from group comparisons and towards individual-level analyses, an essential step on the road to precision medicine (Chamberland et al., 2021; Marquand et al., 2016; Wolfers et al., 2018).

Multivariate approaches that combine structural MRI metrics have been used in a number of promising contexts. At the group level, partial least squares (PLS) analyses and their variants can be used to assess the covariance between multiple metrics (Khedher et al., 2015; Nestor et al., 2002). Other multivariate approaches that can be used in group analyses include principal component analysis (PCA), independent component analysis (ICA) and non-negative matrix factorization (Calhoun et al., 2001; Khedher et al., 2015; Plitman et al., 2020; Yang et al., 2011). At the individual level, inter-regional correlations of multiple metrics can be used to create individual-specific network maps based on morphometric similarity that can then be linked to behavior (Seidlitz et al., 2018). Individualized network maps provide a more comprehensive structural mapping that captures both biological complexity and individual variability because they integrate multiple MRI features (e.g., Vandekar et al., 2016; Whitaker et al., 2016).

However, in this study by Seidlitz et al., (2018), the shared covariance between MRI metrics was not accounted for. This has the potential to bias inferences made from such analyses, as there is significant covariance between many commonly used imaging parameters (Carter et al., 2022; Uddin et al., 2019). Various multivariate approaches that can overcome this issue exist, including multivariate linear regression (Naylor et al., 2014; Young et al., 2010), machine-learning (e.g., Calhoun et al., 2001; Carbonell et al., 2020; Chen et al., 2019; Guberman et al., 2022; Khedher et al., 2015; Yang et al., 2011), and Hotelling’s T^2^ test (Avants et al., 2008; Hotelling, 1947). However, many of these approaches (including multivariate linear regression and machine learning) are computationally expensive and some necessitate making subjective decisions (Alexopoulos, 2010; Gyebnár et al., 2019; Hayasaka et al., 2006; Naylor et al., 2014). The Hotelling’s T^2^ test, a multivariate extension of a two-sample t-test, is a simple yet powerful option for group comparisons (Avants et al., 2008; Hotelling, 1947), but provides little insight at the individual level (Guberman et al., 2022).

Here we propose using the Mahalanobis distance (D2) (Mahalanobis, 1936) for analyzing multimodal MRI metrics. D2 is closely related to Hotelling’s T^2^, but can also provide an individual-level measure of deviation relative to a reference distribution. It is defined as the multivariate distance between a point and a distribution in which covariance between features (i.e., imaging metrics) is accounted for. Initially developed by P. C. Mahalanobis in 1936 to quantify racial similarities based on anthropometric measurements of skulls (Mahalanobis, 1927), D2 can be thought of as a multivariate z-score where the covariance between features is accounted for (Taylor et al., 2020).

The D2 approach has been used extensively in outlier detection, cluster analysis, and classification applications (e.g., Ghorbani, 2019; Kritzman & Li, 2010; Xiang et al., 2008). D2 has also been used in neuroimaging, mainly in the study of neurological disorders, to detect lesions (Gyebnár et al., 2019; Lindemer et al., 2015), or to evaluate the degree of abnormality in the brains of patients relative to controls (Dean et al., 2017; Owen et al., 2021; Taylor et al., 2020), but also to study healthy WM development (Kulikova et al., 2015). Despite promising implementations and its high versatility, D2 has not yet been widely adopted. To facilitate its use, we present here an open-source python-based tool for computing D2 relative to a reference group or within a single individual: the MultiVariate Comparison (MVComp) toolbox. We provide a step-by-step guide to computing D2 using the MVComp tool (https://github.com/neuralabc/mvcomp) for two distinctive scenarios: a) comparisons between a subject and a reference group, and b) within-subject comparisons between voxels (Section 2). Lastly, the results of these example cases are presented (Section 3) and the general approach is discussed (Section 4) (Tremblay et al., 2024).

## 2. Methods

### 2.1 General framework

Since D2 can be defined relative to virtually any reference of matching features, MVComp has been designed to support a wide range of applications. The first step is to define the set of multivariate data that will serve as the reference for computing D2. This choice depends on the hypothesis of interest, which will determine the *Level of Analysis* (Fig. 1). D2 can be computed between different brain regions within an individual (with the individual’s data also serving as the reference) or between an individual and a group, in spatially correspondent regions. In each case, multiple different *Resolutions* of analysis are possible, including voxel-wise and region of interest-(ROI) based comparisons.

**Fig. 1.**
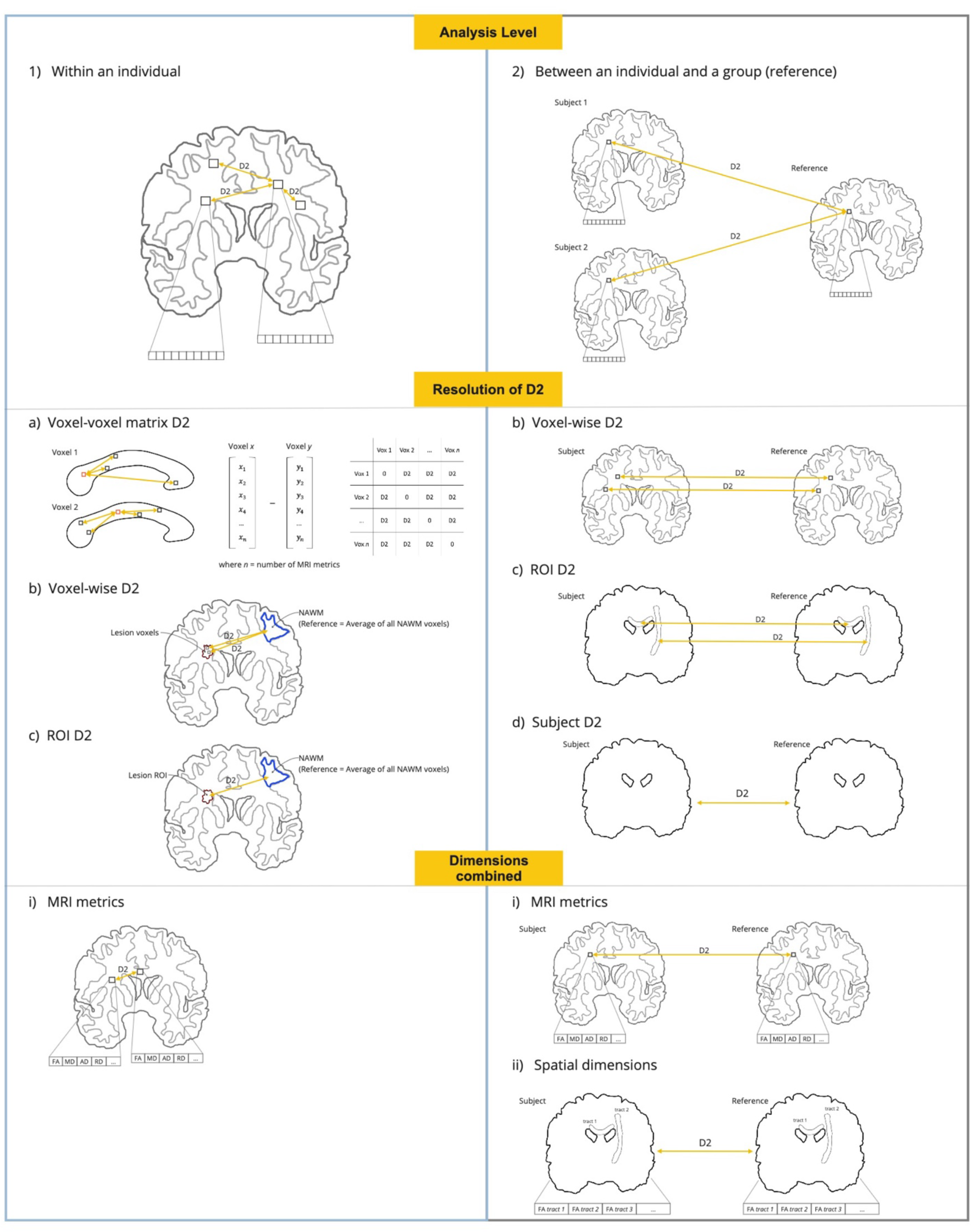
Implementations of the D2 framework in neuroimaging studies. **Analysis level:** (1) Within an individual (left panel, in light blue): D2 can be computed between different voxels or brain regions (e.g., WM tracts) within a single subject. (2) Between an individual and a group (right panel, in light gray): D2 can be computed between a subject and a reference group (e.g., control group). **Resolution of D2:** (a) Voxel-voxel matrix D2: D2 can be computed between each voxel and all other voxels in a mask of analysis, resulting in a D2 matrix of size *n* voxels x *n* voxels (only applicable to analyses within an individual). (b) Voxel-wise D2: A D2 value can be computed at each voxel. (c) ROI D2: In this case, a D2 value is obtained for each WM tract, or other brain region (ROI) defined by the user. (d) Subject D2: A single D2 value can be obtained per subject, resulting in a measure of global brain deviation from the reference (only applicable to analyses between an individual and a group). **Dimensions combined:** (i) MRI metrics: when the dimensions combined through D2 are MRI metrics, the length of the vector of data is the number of metrics. (ii) Spatial dimensions: when WM tracts, or other parcellated brain regions, are combined through D2, the length of the vector of data is equal to the number of WM tracts (only applicable to analyses between an individual and a group; yields a single D2 value per subject).

Lastly, the choice of which dimensions to combine, either MRI-derived metrics or brain regions (e.g., WM tracts), depends on what we want to capture. Combining brain regions within a multivariate measure allows to capture the degree of deviation from a reference even in the presence of high spatial heterogeneity (e.g., Owen et al., 2021; Taylor et al., 2020), while combining features is useful in the presence of mechanistic heterogeneity (i.e, several concomitant underlying biological mechanisms) and when preserving regional specificity is desirable (e.g., Guerrero-Gonzalez et al., 2022; Gyebnár et al., 2019; Lindemer et al., 2015). See Fig. 1. for a comprehensive view of the possible combinations of levels of analysis, resolutions and with different dimensions combined.

To illustrate the flexibility of the D2 approach, we present a few examples:

#### 2.1.1 Comparisons between an individual and a group (reference)

**Example 1:** Computing a voxel-wise D2 map for each individual

**Data**: Diffusion MRI (dMRI) data in several subjects

**Level of Analysis:** Between an individual and a group (Fig. 1 right panel)

**Feature Resolution:** Voxel-wise D2 (in all WM voxels) (Fig. 1b)

**Dimensions combined:** dMRI-derived metric maps (Fig. 1i)

In this example the reference would be defined as the voxel-wise group average for each dMRI-derived metric (m_1_, m_2_, m*_n_,* where *n* is the number of metrics) and D2 is computed by comparing the feature values in each voxel of an individual to the corresponding voxel in the reference (see Fig. 2a-c). The resulting D2 maps can then be entered into second-level analyses to, for example, identify brain-behavior associations. If two groups are being analyzed (e.g., patients vs controls), the control group could be used as the reference and D2 values computed between each patient and the reference would represent voxel-wise multivariate distance from controls.

**Fig. 2.**
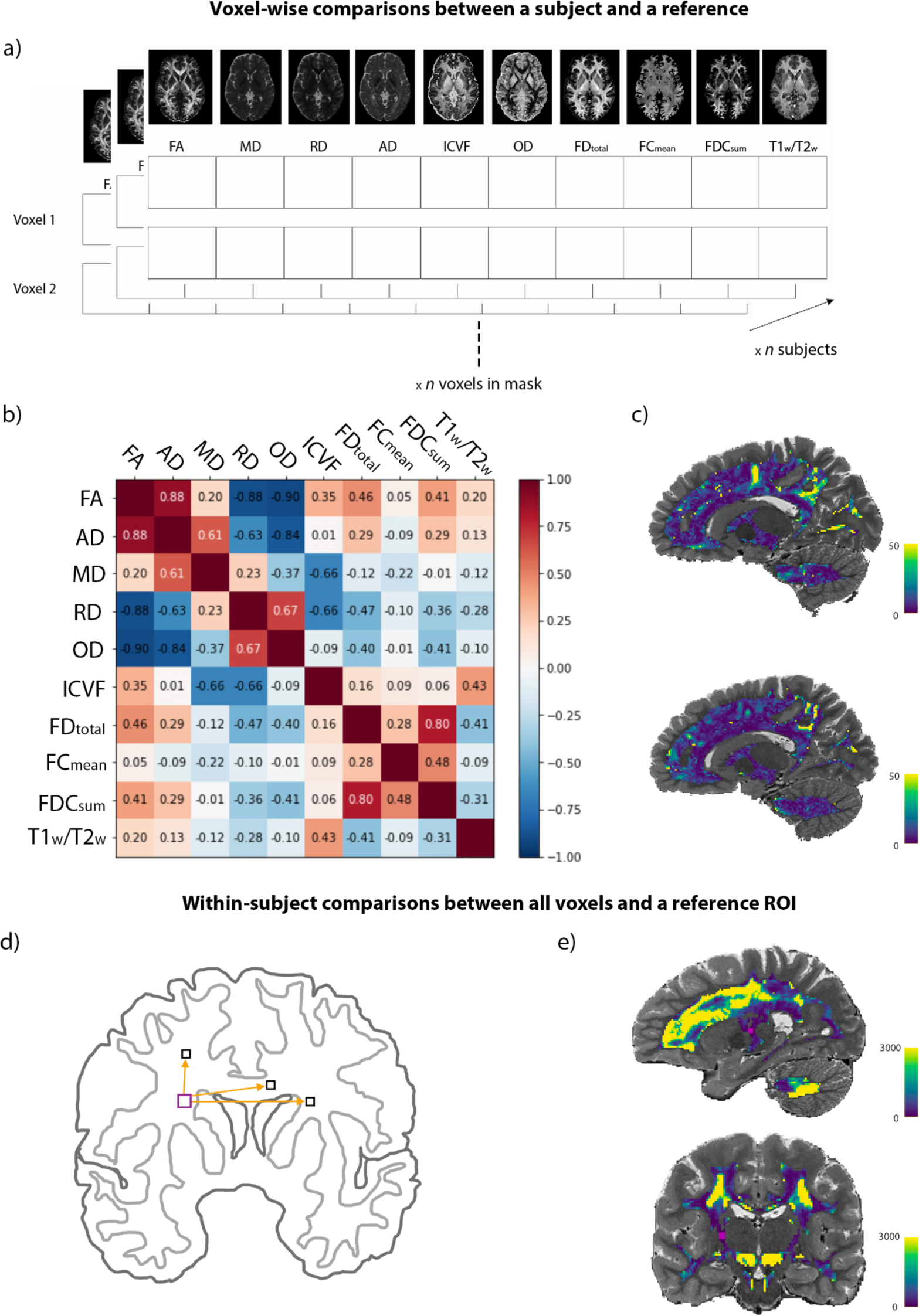
D2 workflow. **Voxel-wise comparisons between a subject and a reference.** (a) The multivariate space is illustrated here. In this example, we have a vector of 10 dMRI metrics at each WM voxel for each subject. (b) The covariance matrix is computed from the reference feature matrix of shape *n* voxels in WM x *n* features. The plot shows the amount of correlation between features in the reference sample (i.e., the whole group). (c) Voxel-wise D2 maps in two example subjects, where bright yellow represents areas of greater deviation from the reference population. Distinct patterns can be seen in the two subjects. Note that the leave-one-subject-out approach was used so that the data of the subject under evaluation was not included in the group mean (i.e., reference) and covariance matrix prior to D2 calculation. **Within-subject comparisons between all WM voxels and a reference ROI.** (d) Schematic representation of the multivariate comparisons showing that D2 was computed between all WM voxels and a ROI of 24 voxels in the corticospinal tract (CST). (e) D2 map showing the multivariate distance between all WM voxels and the CST ROI (in pink). *Data used for these examples will be presented in section 2.7.

**Example 2:** Computing a single D2 score per individual

**Data**: dMRI data in several subjects

**Level of Analysis:** Between an individual and a group (Fig. 1 right panel)

**Feature Resolution:** Subject D2 (Fig. 1d)

**Dimensions combined:** WM tracts (spatial dimensions) (Fig. 1ii)

A single MRI metric can also be used and combined across multiple ROIs (e.g., mean FA in pre-defined WM tracts). The reference is defined as the group mean of each tract (*m_1_, m_2_, m_n_,* where *n* is the number of tracts) and a single D2 value is computed for each individual. In this case, D2 represents a measure of how different an individual’s WM microstructure is relative to a reference, across multiple tracts. This application is not demonstrated in the present article but it has been used by others (e.g., Owen et al., 2021; Taylor et al., 2020) and can be implemented using MVComp.

To ensure that each subject’s data will not bias their D2 values in single sample designs (i.e., where the entire sample is used as a reference) and to allow the evaluation of controls in two-sample designs, a leave-one-subject-out approach is also possible. In this way, the subject under evaluation is excluded from the group mean (reference) and covariance matrix prior to calculating D2.

#### 2.1.2 Comparisons within an individual

**Example 3:** Computing D2 between lesion voxels and normal appearing WM (NAWM)

**Data**: dMRI data in one subject

**Level of Analysis:** within an individual (Fig. 1 left panel)

**Feature Resolution:** voxel-wise (in lesion voxels) (Fig. 1b)

**Feature Dimensions:** dMRI-derived metric maps (Fig. 1i)

Here, the level of analysis is within-subject, the dimensions combined are multiple dMRI-derived metrics in each voxel, and the reference is the average of all voxels within a region of interest (ROI) for each metric. To investigate the distance between WM lesions and NAWM, the reference would be defined as the average of all NAWM voxels (*m_1_, m_2_, m_n_,* where *n* is the number of metrics) and D2 would be computed for each voxel classified as a lesion. Alternatively, the *resolution* could be ROI-wise, if the user deems a single D2 value per lesion sufficient. This within-subject approach can also be used as a measure of similarity by computing D2 between all WM voxels and a reference ROI in a specific tract (e.g., voxels in the cortico-spinal tract, as in Fig. 2d). Voxels within the same WM tract as the reference ROI are likely to have lower D2 values (indicating higher similarity) than voxels in other tracts or in areas of crossing fibers (Fig. 2e).

**Example 4:** Computing D2 between each voxel and all other voxels in a mask

**Data**: dMRI data in one subject

**Level of Analysis:** within an individual (Fig. 1 left panel)

**Feature Resolution:** Voxel-voxel D2 matrix (Fig. 1a)

**Feature Dimensions:** dMRI-derived metric maps (Fig. 1i)

D2 can be calculated between every pair of voxels (voxel *x* − voxel *y*) within a mask of analysis to compute a voxel-voxel D2 matrix (see Fig. 1a). In this case, the reference for computing the covariance matrix would be the data in all voxels contained in the mask.

### 2.2 Data preparation

In all cases, data for all subjects should be preprocessed and all MRI metrics of interest computed and transformed to bring them into the same voxel space. If instead of voxel-wise comparisons the user is interested in performing ROI-based comparisons, summary metrics should be calculated for each region of interest (e.g., mean FA in each WM tract of interest) for each subject. Masks should also be generated to restrict analyses to chosen regions (e.g., WM) and these should also be transformed into the same space. Masks can be binary or thresholded at a later step within MVComp.

### 2.3 Computing the reference mean and covariance matrix

In the case of analyses between subject(s) and a reference (Fig. 1 right panel), the reference mean and covariance matrix are derived either from multiple features (Fig. 1i) or multiple ROIs (Fig. 1ii) in another group (e.g., control group). The comparison can also be between each individual and the mean of all other individuals if only a single group is available. In the case of analyses within an individual (Fig. 1 left panel), multiple features can be compared between voxels (e.g., Fig. 1 a-b) or between ROIs (e.g., Fig. 1c).

#### 2.3.1 Comparisons between an individual and a group (reference) Combining MRI metrics

For this application, the group average of each metric must be computed from the reference group (`mvcomp.compute_average` can be used to perform this task). The `mvcomp.feature_list` function can then be used to create a list of feature names and a list of the full paths of the average maps that were created with the `compute_average` function. The `mvcomp.feature_gen` function extracts the feature matrix from a set of input images. Run on the reference group mean images with a provided mask, it returns the feature matrix (`m_f_mat` of shape *n* voxels in the mask x *n* features), a mask vector (`mat_mask` of shape *n* voxels) and a nibabel object of the mask (`mask_img`). The mask array contains zeros at voxels where values are *nan* or *inf* for at least one of the reference average maps in addition to the voxels below the threshold set for the mask. The `mvcomp.norm_covar_inv` function is then used to compute the covariance matrix (`s`) and its pseudoinverse (`pinv_s`) from the reference feature and mask matrices (`m_f_mat` and `mat_mask)`. The `mvcomp.correlation_fig` function can be used to generate a correlation matrix from the covariance matrix (`s`), which is informative to verify if expected relationships between features are present.

A leave-one-out approach (where the individual to be compared to the reference is left out of the average) is preferred in cases where the individual subject is also a member of the reference group. This functionality is directly available in the model comparison function (`model_comp`). If the leave-one-out approach is used, it is not necessary to compute the group average nor to use the `mvcomp.feature_gen` and `mvcomp.norm_covar_inv` functions since the average and covariance matrix will be computed within the `model_comp` function from a group that excludes the subject for which D2 is being computed.

##### Combining spatial dimensions

The reference mean values (e.g., reference group mean FA in each WM tract) and covariance matrix are computed within the `spatial_mvcomp` function described in detail below. See Owen et al., 2020; Taylor et al., 2020 for example applications of this implementation.

#### 2.3.2 Comparisons within an individual Voxel-wise D2 resolution

In the case of comparisons within a single subject, one of the possible applications is to compute D2 between specific ROIs. If the reference ROI is a set of NAWM voxels, the covariance matrix will be computed based on all voxels within that ROI in that subject. The path of the images (i.e., one image per metric) can be provided to the `feature_gen` function, along with the ROI mask, to create the reference feature matrix (`m_f_mat`) and mask vector (`mat_mask`). The `mvcomp.norm_covar_inv` function is then used to compute the covariance matrix (`s`) and its pseudoinverse (`pinv_s`) from the feature and mask matrices. The `mvcomp.correlation_fig` function can again be used to visualize relationships between metrics.

##### Voxel-voxel matrix D2 resolution

For this approach, the covariance matrix is computed from a feature matrix that includes all voxels in the mask of analysis. For instance, if we are interested in computing D2 between each voxel and all other voxels in the whole WM, the covariance matrix is based on all WM voxels. Therefore, the matrix provided to the `norm_covar_inv` function will be of shape *n* voxels in the mask x *n* features.

### 2.4 Computing D2

Once the mean of the reference and the covariance matrix have been computed and the data for comparisons has been prepared, the D2 computation can be performed. Depending on the *resolution* of D2, this computation may be repeated several times (i.e., between every pair of voxels or once for each voxel or each ROI; Fig. 1a-c), or it may only be done once if the user is interested in obtaining a single individualized score of deviation from a group (Fig. 1d). The MVComp tool contains functions to easily compute D2 for each of these applications, according to this equation:

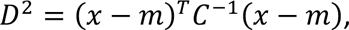

where % is the vector of data for one observation (e.g., one subject), *m* is the vector of averages of all observations for each independent variable (e.g., MRI metrics), and (*C^-1^* is the inverse of the covariance matrix.

#### 2.4.1 Comparisons between an individual and a group (reference) Combining MRI metrics

The `mvcomp.model_comp` function allows the calculation of voxel-wise D2 between each subject contained in the provided subject_ids list and the reference (group average) (Fig. 1 right panel; b). The user should specify the directories and suffix of the subjects’ features and of the reference images (feature_in_dir, model_dir, suffix_name_comp and suffix_name_model), the mask of analysis (mask_f) and a threshold if the mask is not binary (mask_threshold). If subjects or features are to be excluded at this point, they can be specified with the exclude_subject_ids and the feat_sub options, respectively. If the user wishes to use the leave-one-out approach, the exclude_comp_from_mean_cov option should be set to True. If this option is set to True, the mean (reference) and pinv_s are computed for each subject comparison, excluding the subject being compared before computing its D2. Therefore, it is not necessary to specify the directory of the reference (model_dir) in this application. The `model_comp` function yields a matrix containing the D2 data of all subjects (of size number of voxels x number of subjects). To obtain a D2 map (in nifti format) for each subject, the dist_plot function can then be used. The function also outputs a mean D2 map of all subjects and a histogram of all D2 values.

##### Combining spatial dimensions

The `mvcomp.spatial_mvcomp` function is used to compute a D2 score between each subject and the reference computed from all subjects (Fig. 1ii). A matrix containing the data (e.g., mean FA in each WM tract) of all subjects (*n* subjects x *n* tracts) should be provided to the function. The `spatial_mvcomp` function returns a vector with a single D2 value per subject, reflecting the subject’s individualized score of deviation from the group. As in `model_comp`, setting the `exclude_comp_from_mean_cov` to True leaves out the current subject when computing the mean and covariance.

#### 2.4.2 Comparisons within an individual Voxel-wise D2 resolution

The `mah_dist_mat_2_roi` function is used to compute voxel-wise D2 between all voxels within a mask and a specific ROI (Fig. 1 left panel; b). Here, in addition to the feature matrix containing the data for the voxels to be evaluated (*n* voxels in the mask x *n* features), the user will need to provide a vector of data for the reference ROI (i.e., mean across voxels in the ROI for each metric) and the inverse of the covariance matrix (`pinv_s`).

##### Voxel-voxel matrix D2 resolution

The `voxel2voxel_dist` function is used to compute D2 between each voxel and all other voxels in a mask (Fig. 1 left panel; a). This yields a symmetric 2-D matrix of size *n* voxels x *n* voxels containing D2 values between each pair of voxels.

### 2.5 Statistical analysis

Once D2 values are computed, second-level statistical analyses can be used to assess group differences and longitudinal trajectories, to explore relationships between D2 and behavior. Machine learning techniques can also be used to reduce dimensionality and produce network maps based on (dis)similarity.

#### 2.5.1 Comparisons between an individual and a group (reference)

For group comparisons, a two-samples t-test can be performed on D2 values (e.g., D2 values in patients vs D2 in controls), which would be equivalent to performing a Hotelling’s T^2^ test on raw metrics (i.e., without computing D2). Alternatively, a statistical method such as the Bhattacharyya coefficient can be used to estimate the degree of overlap between the distribution of each group, where less overlap indicates a higher probability that the groups differ, as in (Dean et al., 2017). However, such group analyses are likely to average out interindividual variability and may be problematic when heterogeneity is high (Guberman et al., 2022). Wilk’s criterion is another approach that can be used to define abnormality based on a calculated critical value that accounts for normative sample size, number of features, and multiple comparisons (Guerrero-Gonzalez et al., 2022; Gyebnár et al., 2019; Wilks, 1963).

#### 2.5.2 Comparisons within an individual

In within-subjects analyses, clustering approaches can be applied to the voxel-voxel matrix D2 to partition brain voxels into networks or other parcellations. Changes in D2, either from the group or subject-level, can also be assessed through longitudinal analyses, to investigate WM damage progression or brain maturation for instance (e.g., Kulikova et al., 2015; Lindemer et al., 2015). D2, or changes in D2, can also be related to behavioral outcomes (e.g., cognitive score, performance on a skill test, or symptom severity) in the same way one would with univariate measures of fractional anisotropy for instance (Dean et al., 2017; Owen et al., 2021; Taylor et al., 2020).

### 2.6 Determining feature importance

D2 summarizes the amount of deviation from a reference, based on several metrics or brain regions, into a single score. This yields a useful metric to easily quantify *abnormalities*, whether due to pathology or to exceptional abilities such as musical skills. However, when summarizing several features into a single score, we lose specificity. To help address this limitation, it is possible to extract the contribution of each feature to the multivariate distances (D2) using functions of the MVComp tool to recover biological or spatial specificity.

#### 2.6.1 Comparisons between an individual and a group (reference) Combining MRI metrics

If the user is interested in understanding the physiological mechanisms underlying microstructural deviations in a region of interest (e.g., voxels where D2 is high), the `return_raw` option of the `mvcomp.model_comp` function can be used. This allows the extraction of each metrics’ weight in D2. If `return_raw` is set to True, the function returns a 3D array of size (number of voxels) x (number of metrics) x (number of subjects) that contains the voxel-wise distances for each feature and each subject. A flattened mask of the region of interest (e.g., a region of high D2) can then be applied to select voxels from the 3D array. The distances can be summarized across voxels and/or subjects to obtain a % contribution to D2 for each MRI metric within that region.

##### Combining spatial dimensions

The `return_raw` option is also available in the `spatial_mvcomp` function. If set to True, a 2D array of size (number of subjects) x (number of tracts) containing the distances between every subject’s tract and the mean tract values is returned. These *raw* distances provide information regarding the contribution of each WM tract to D2, which gives insights on the localization of greatest deviation for each subject.

#### 2.6.2 Comparisons within an individual Voxel-wise D2 resolution

The return_raw option of the mah_dist_mat_2_roi function can be used to extract features’ contributions. In this case, the distances between features in all voxels being compared and feature values in the ROI are returned. The output will be of shape (number of voxels) x (number of metrics).

### 2.7 Experiments

#### 2.7.1 Data Description

We computed 10 microstructural features for 1001 subjects from the Human Connectome Project S1200 data release (Van Essen et al., 2013) for these experiments. DWI, T1- and T2-weighted data were acquired using a custom-made Siemens Connectom Skyra 3 Tesla scanner with a 32-channel head coil. The DWI data (TE/TR=89.5/5520 ms, FOV=210×180 mm) were multi-shell with b-values of 1000, 2000 and 3000 s/mm^2^ and a 1.25 mm isotropic resolution, 90 uniformly distributed directions, and 6 b=0 volumes. T1-w data was acquired with a 3D-MPRAGE sequence and T2w images with a 3D T2-SPACE sequence, both with a 0.7mm isotropic resolution (T1w: 0.7 mm iso, TI/TE/TR=1000/2.14/2400 ms, FOV=224×224 mm; T2w: 0.7 mm iso, TE/TR=565/3200 ms, FOV=224×224 mm). Anatomical scans were acquired during the first session, and DWI data were acquired during the fourth session. More details on the acquisitions can be found at: https://www.humanconnectome.org/hcp-protocols-ya-3t-imaging. The imaging data of 1065 young healthy adults, those who had undergone T1w, T2w and diffusion-weighted imaging, were preprocessed. The data of 64 participants were excluded due to poor cerebellar coverage.

#### 2.7.2 Preprocessing Diffusion Tensor Imaging

The minimally preprocessed HCP data was used (Glasser et al., 2013; Van Essen et al., 2013). The minimal preprocessing pipeline for DWI data includes intensity normalization of the b_0_ images, eddy current and susceptibility-induced distortions correction, using DWI volumes of opposite phase-encoding directions, motion correction and gradient nonlinearity correction. DWI data were registered to native structural space (T1w image), using a rigid transform computed from the mean b_0_ image, and diffusion gradient vectors (bvecs) were rotated accordingly.

Most subsequent processing steps were performed using the MRtrix3 toolbox (Tournier et al., 2019). The minimally preprocessed DWI data was converted to the mif format, with the bvals and bvecs files embedded, after which a bias field correction was performed using the ANTs algorithm (N4) of the dwibiascorrect function of MRtrix3 (Tustison et al., 2010). The tensor was computed on the bias field-corrected DWI data (using *dwi2tensor*) and DTI metrics were then calculated (FA, MD, AD and RD) using *tensor2metric* (Basser et al., 1994a, 1994b; Veraart et al., 2013).

##### Multi-tissue Multi-shell Constrained Spherical Deconvolution

The multi-tissue Constrained Spherical Deconvolution (CSD) was performed following the fixel-based analysis (FBA) workflow <u>(Tournier et al., 2019)</u>. The T1-w images were segmented using the *5ttgen* FSL function of MRtrix3, which uses the FAST algorithm (Patenaude et al., 2011; R. E. Smith et al., 2012; S. M. Smith, 2002; S. M. Smith et al., 2004; Y. Zhang et al., 2001). Response functions for each tissue type were then computed from the minimally preprocessed DWI data (without bias field correction) and the five-tissue-type (5tt) image using the *dwi2response* function (msmt_5tt algorithm) (Jeurissen et al., 2014). The bias-uncorrected DWI data was used because bias field correction is performed at a later step in the FBA pipeline (Raffelt, Tournier, et al., 2017). The WM, GM and CSF response functions were then averaged across all participants, resulting in a single response function for each of the three tissue types. Multi-shell multi-tissue CSD was then performed based on the response functions to obtain an estimation of orientation distribution functions (ODFs) for each tissue type (Jeurissen et al., 2014). This step is performed using the *dwi2fod msmt_csd* function of MRtrix3 within a brain mask (i.e., nodif_brain_mask.nii.gz). Bias field correction and global intensity normalization, which normalizes signal amplitudes to make subjects comparable, were then performed on the ODFs, using the *mtnormalise* function in MRtrix3 (Dhollander et al., 2021; Raffelt, Dhollander, et al., 2017).

##### Registration

In order to optimize the alignment of WM as well as gray matter, multi-contrast registration was performed. Population templates were generated from the WM, GM and CSF FODs and the “nodif” brain masks of a subset of 200 participants using the *population_template* function of MRtrix3 (with regularization parameters: nl_update_smooth= 1.0 and nl_disp_smooth= 0.75), resulting in a group template for each of the three tissue types (Tournier et al., 2019).

Subject-to-template warps were computed using *mrregister* in MRtrix3 with the same regularization parameters and warps were then applied to the brain masks, WM FODs, DTI metrics (i.e., FA, MD, AD and RD), T1w, and T2w images using *mrtransform* (Raffelt et al., 2011). T1w and T2w images were kept in native resolution (0.7mm) and the ratio of T1w/T2w was calculated to produce a myelin map (Glasser & Essen, 2011). WM FODs were transformed but not reoriented at this step, which aligns the voxels of the images but not the *fixels (“fibre bundle elements”)*. A template mask was computed as the intersection of all warped brain masks (*mrmath min* function). This template mask includes only the voxels that contain data in all subjects. The WM volumes of the five-tissue-type (5tt) 4-D images were also warped to the group template space since these are then used to generate a WM mask for analyses.

##### Computing fixel metrics

The WM FOD template was segmented to generate a *fixel* mask using the *fod2fixel* function (Raffelt et al., 2012; R. E. Smith et al., 2013). This mask determines the fiber bundle elements (i.e., *fixels*), within each voxel of the template mask, that will be considered for subsequent analyses. *Fixel* segmentation was then performed from the WM FODs of each subject using the *fod2fixel* function, which also yields the apparent fibre density (FD) metric. The *fixelreorient* and *fixelcorrespondence* functions were then used to ensure subjects’ fixels map onto the fixel mask (Tournier et al., 2019).

The fibre bundle cross-section (FC) metric was then computed from the warps generated during registration (using the *warp2metric* function) as FC is a measure of how much a fiber bundle has to be expanded/contracted for it to fit the fiber bundles of the *fixel* template. Lastly, a combined metric, fibre density and cross-section (FDC), representing a fibre bundle’s total capacity to carry information, was computed as the product of FD and FC.

##### Transforming fixel metrics into voxel space

In order to integrate all metrics into the same multi-modal model, *fixel* metric maps were transformed into voxel-wise maps. As a voxel aggregate of fiber density, we chose to use the l=0 term of the WM FOD spherical harmonic expansion (i.e., 1^st^ volume of the WM FOD, which is equal to the sum of FOD lobe integrals) to obtain a measure of the total fibre density (FD_total_) per voxel. This was shown to result in more reproducible estimates than when deriving this measure from fiber specific FD (i.e., by summing the FD *fixel* metric) (Calamante et al., 2015). The FOD l=0 term was scaled by the spherical harmonic basis factor (by multiplying the intensity value at each voxel by the square root of 4π).

For the fiber cross-section voxel aggregate measure, we opted for computing the mean of FC, weighed by FD (using the mean option of the *fixel2voxel* function). We thus obtained the typical expansion/contraction necessary to align fiber bundles in a voxel to the *fixels* in the template.

Lastly, the voxel-wise sum of FDC, reflecting the total information-carrying capacity at each voxel, was computed using the *fixel2voxel sum* option.

##### NODDI metrics

Bias field corrected DWI data was fitted to the neurite orientation dispersion and density imaging (NODDI) model using the python implementation of Accelerated Microstructure Imaging via Convex Optimization (AMICO) (Daducci et al., 2015; H. Zhang et al., 2012). First, small variations in b values were removed by assigning the closest target bval (0, 1000, 2000 or 3000) to each value of the bvals file. This is to prevent the fitting algorithm from interpreting every slightly different bval as a different diffusion shell. A diffusion gradient scheme file is then created from the bvecs, and the new bvals file. The response functions are computed for all compartments and fitting is then performed on the unbiased DWI volumes, within the non-diffusion weighted brain mask (nodif_brain_mask.nii.gz). The resulting parameters obtained are: the intracellular volume fraction (ICVF, also referred to as neurite density), the isotropic volume fraction (ISOVF), and the orientation dispersion index (OD). In this study, we will use ICVF and OD.

##### Generating masks for analyses

The maps of each of the 10 metrics of interest (FA, AD, RD, MD, T1w/T2w, FDtotal, FCmean, FDCsum, ICVF and OD) were then averaged across all subjects. These average maps served as the reference. A WM mask was created by computing the group average of the corresponding volume of the T1 5tt image (volume 2). A threshold of 0.99 was applied within the MVComp toolbox’s functions.

#### 2.7.3 Experiment 1: Comparisons between an individual and a group (reference)

Here, we present an example case of using D2 in a large sample from the HCP dataset to quantify voxel-wise microstructural differences in WM according to several dMRI metrics. Since the dataset used in this study contains the data of healthy young adults, a relatively homogeneous population, the entire sample was set as the reference and the leave-one-out approach was used to exclude the subject under evaluation. The analysis was restricted to the corpus callosum (CC). Voxel-wise D2 from 10 microstructural features was computed in the CC for each subject, yielding a D2 matrix of 1001 subjects X 2845 voxels. The D2 values represent voxel-wise microstructural differences in an individual’s CC relative to the group average, while accounting for the covariance between features. Large D2 scores in a voxel indicate greater deviation from the group average, whereas scores closer to 0 indicate lower distance (i.e., more typical microstructure).

Past literature on CC neuroanatomy shows several segments that are distributed along the anterior to posterior axis, where each segment is defined by common microstructural properties and/or connectivity profiles (Aboitiz et al., 1992; Chao et al., 2009; Hofer & Frahm, 2006). We therefore hypothesized that these segments could be extracted via clustering, an unsupervised machine learning technique, of D2 values in the CC. We performed k-means clustering on the D2 matrix, setting the number of clusters to 9 based on literature on CC topography (Aboitiz et al., 1992; Chao et al., 2009; Hofer & Frahm, 2006). Prior to clustering, we applied z-score and power transformation on the D2 matrix to achieve gaussian distributions of the standardized scores. Due to the large number of datapoints and potential effects of partial voluming, we observed several outliers in D2 maps of several subjects. We therefore excluded participants with at least 50 voxels that were deemed as outliers (i.e. exceeded a threshold of 5 standard deviations from the voxel mean D2). This yielded a final sample of 723 participants. Final visualization was done using BrainNet Viewer (http://www.nitrc.org/projects/bnv/).

#### 2.7.4 Experiment 2: Comparisons within an individual

The within-subject approach allows the computation of voxel-voxel D2 in a single individual from multiple microstructural features. Here, D2 was calculated between each voxel and every other voxel in a subject’s CC, while accounting for the covariance between the 10 microstructural features. All voxels within the CC of that subject were used to compute the covariance matrix and this same covariance matrix was used in the D2 calculation of every voxel. The resulting D2 matrix is a 2845 voxel X 2845 voxel dense matrix representing the distance between each voxel and every other voxel in the CC (Fig. 4a-b). We standardized the matrix to z-scores and applied Principal component analysis (PCA) to reduce the matrix dimensionality (Fig. 4c). We then extracted the contributions of each metric to D2 within the voxels with the largest and the lowest scores on the first principal component (Fig. 4d-f).

## 3. Results

### 3.1 Experiment 1: Comparisons between an individual and a group (reference)

For this experiment, D2 was computed voxel-wise in the CC between each subject and a reference consisting in all other subjects (Fig. 3a-b). K-means clustering was applied to the D2 matrix of size (subjects) X (voxels). We observed that the 9 clusters were distributed along the anterior-posterior axis, in accordance with past evidence on CC microstructure and connectivity (Aboitiz et al., 1992; Chao et al., 2009; Hofer & Frahm, 2006). Fig. 3c shows the clusters identified via k-means and Fig. 3d shows the topography expected according to literature. The genu of the CC was clustered into 3 segments, while the midbody displays 2 segments. The splenium was divided into 4 segments (with one segment positioned on the isthmus).

**Fig. 3.**
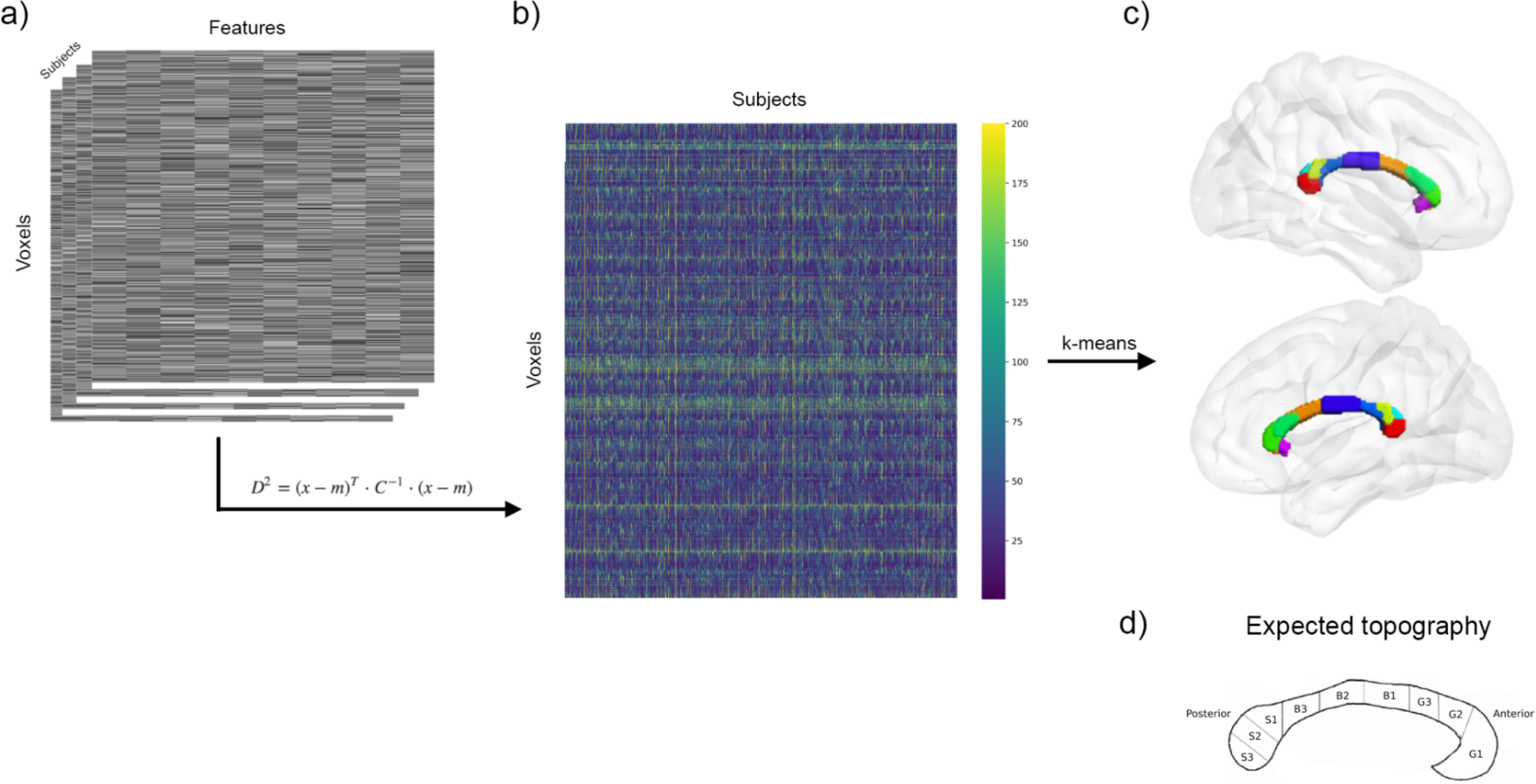
Voxel-wise comparisons between each subject and the reference. (a) Voxel-wise D2 is calculated between the reference (group average of the whole sample, except the subject under evaluation) and each subject’s data (feature (10) X voxel (2845) matrix), in voxels of the corpus callosum (CC). (b) This results in a D2 matrix of size subject (723 after exclusion of outliers) X voxel (2845) containing the multivariate distance between a subject’s data and the reference at each CC voxel. (c) Applying k-means clustering to the D2 matrix, voxels of the CC were partitioned into 9 clusters distributed along the anterior-posterior axis, in close accordance with known topography of the CC as seen in (d). (d) Schematic representation of CC topography based on literature (Aboitiz et al., 1992; Chao et al., 2009; Hofer & Frahm, 2006).

### 3.2 Experiment 2: Comparisons within an individual

For the within-subject experiment, D2 was computed between all voxel pairs in the CC of a single individual, yielding a voxel X voxel D2 matrix (Fig. 4a-b). PCA was applied to the D2 matrix. Fig. 4c shows the first 10 principal components (PCs). We then extracted the contributions (i.e., loadings) of each metric to D2 within the voxels with the largest and the lowest scores on the first principal component. The first PC explained 95% of the variance in the voxel X voxel dense D2 matrix. The highest and the lowest PC1 scores were in the genu and in the midbody of the CC, respectively (Fig. 4d). In the voxel with the largest value on PC1, the fibre density and cross-section metric (sumFDC) contributed most to D2, while mean diffusivity (MD) contributed the least (Fig. 4f). On the other hand, in the voxel with the lowest score on PC1, all microstructural features had nearly equal contributions to D2, indicating minimal variability in this voxel (Fig. 4e).

**Fig. 4.**
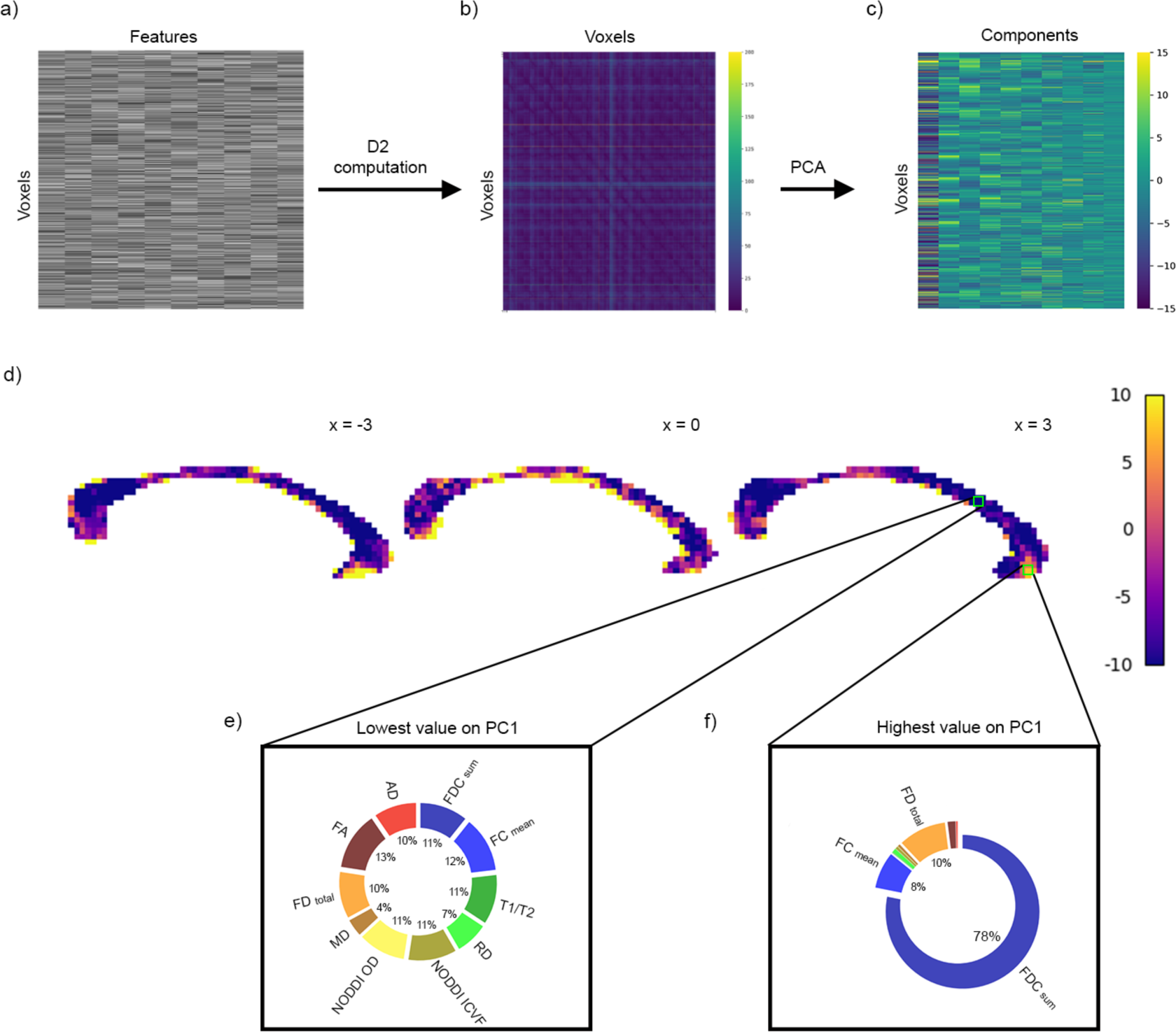
Within-subject voxel-voxel comparisons. D2 was computed between all voxel pairs from the (a) (features) x (voxels in the CC) matrix of a subject. (b) A voxel x voxel D2 matrix was generated. (c) PCA was then applied to the D2 matrix. The PCA matrix shows the first 10 principal components. (d) Voxels with the highest and lowest score on PC1 are shown. PC1 scores were scaled between −10 and 10 to facilitate visualization. (e) In the voxel with the lowest value on PC1, located in the midbody of the CC, all metrics had approximately equal contribution to D2. (f) SumFDC contributed most to D2 in the voxel with the highest PC1 score, located in the genu of the CC.

## 4. Discussion

In the present study, we introduced the MVComp tool, a set of python-based functions that can be used to compute the Mahalanobis distance (D2) for a wide range of neuroimaging applications. At the group-level, MVComp allows the calculation of a score that quantifies how different the brain structure of an individual is from a reference group. The MVComp tool provides a versatile framework that can be used to answer various research questions, from quantifying the degree of abnormality relative to a control group in individuals with a pathology, to exploring interindividual variability in healthy cohorts. At the subject level, D2 can be used to assess differences between regions of interest or to compute a measure of similarity that can then be used for subsequent analyses (e.g., graph theory/network analyses). Lastly, D2 can combine multiple MRI metrics in the same spatial locations, or it can combine a single metric across several brain regions.

Our approach allows the integration of several variables while accounting for the relationships between these variables. Several biological properties influence the same neuroimaging metric and multiple neuroimaging metrics indirectly reflect a similar underlying physiological property. This overlap means that accounting for covariance between metrics is essential. It also means that using a single neuroimaging metric, or metrics stemming from a single model, offers limited potential for interpretation and is biased by the set of assumptions of the chosen model (e.g., some models assume fixed compartment diffusivities while others attempt to estimate them) (Novikov et al., 2018). Similarly, integrating the assessment of multiple brain regions may map better onto behavior (e.g., cognition or disease severity) than assessing each region separately. Here, again the relationships between variables should be accounted for as observations are not completely independent from each other (i.e., in the same individual, there is likely a great amount of covariance between FA in different voxels or in different WM tracts). While some multivariate frameworks have been implemented in the neuroimaging field, several of them are either applicable at the group level or at the subject level (Alexander-Bloch et al., 2013; Hotelling, 1947; Marquand et al., 2016; Seidlitz et al., 2018), and do not extend from one level to another. Moreover, several multivariate approaches are complicated to implement and computationally expensive which limits their accessibility (Alexopoulos, 2010; Gyebnár et al., 2019; Hayasaka et al., 2006). The D2 framework, on the other hand, is highly versatile and the open-source MVComp toolbox we propose makes implementation accessible for assessing various research questions (see Fig. 1).

One of the novelties of this work is that it provides the option to extract the contributions of all features within the D2 measure. This addresses one of the main limitations of typical multivariate frameworks, allowing researchers to develop more mechanistic interpretations. In previous work using the D2 approach, the loadings (or weights) of the elements combined in the multivariate measure (i.e., either WM tracts or MRI metrics) were not extracted, which has been a significant limitation (Dean et al., 2017). Characterizing the extent by which each feature contributes to D2 can provide important insights into the physiological underpinnings of the differences observed and/or their localization. To our knowledge, MVComp is the only available toolbox for computing D2 on imaging data. In this paper, we detailed the usage of MVComp through 4 example cases (see Supplementary material) covering a wide range of applications and presented the results of 2 experiments.

### D2 reflects the underlying microstructure of WM

To provide specific examples of how MVComp can be used, the D2 framework was applied to the assessment of WM microstructure. We found the approach to be particularly suitable for the study of WM because of the number of modeling methods available for dMRI data. However, it is important to note that other types of tissues and imaging techniques can also be used within the MVComp framework. By applying K-Means clustering to D2 in the corpus callosum, we observed a clear segmentation along the anterior-posterior axis (Fig. 3), consistent with known topography from ex-vivo anatomical studies and tractography-based connectivity (Aboitiz et al., 1992; Chao et al., 2009; Hofer & Frahm, 2006). This high correspondence between clustered D2 and previously described CC topography suggests that the microstructural score obtained by combining several WM neuroimaging metrics through D2 provides a useful index of microstructure.

At the individual level, D2 can capture the amount of (dis)similarity between voxels and, through the extraction of features’ contributions (i.e., loadings), the specific microstructural properties underlying regional differences can be inferred. For example, in our within-subject experiment (Fig. 4) we found high spatial heterogeneity in the relative contributions of different features to D2. The voxel with the highest loading on the first latent component (PC1) was primarily dominated by one metric (sumFDC) while the voxel with the lowest loading was characterized by similar weightings across all features. In the voxel with the highest PC1 score, sumFDC (combined metric of fiber cross-section and density, indicative of the amount of information-carrying capacity) contributed most to D2, meaning sumFDC had higher variability across CC voxels than other metrics. This is consistent with the known microstructural properties of the CC, which shows regional variations in densities of fibers of different sizes along the CC (Aboitiz et al., 1992; Hofer & Frahm, 2006). Further, given that the CC is composed of tightly packed fiber tracts, MD would likely be very low in all those CC voxels (i.e., low variability), which would explain its low contribution. Overall, this supports the relevance of D2 in assessing variability in WM microstructure properties and showcases the use of the features contribution option (i.e, `return_raw`) included in MVComp.

### D2 in the study of pathologies

Given the complexity of underlying pathological changes in various brain conditions, multiparametric approaches are a promising avenue to capture the combination of multiple changes in brain properties (Dean et al., 2017; Guberman et al., 2022; Guerrero-Gonzalez et al., 2022; Iturria-Medina et al., 2017; Owen et al., 2021; Taylor et al., 2020). For instance, D2 incorporating fractional anisotropy (FA) in multiple WM tracts in epileptic patients was found to show stronger associations with epilepsy duration than any univariate measure (e.g., mean FA in a single WM tract) (Owen et al., 2021). Another study reported better performance using D2 encompassing FA in several WM tracts, vs using FA in a single tract, in discriminating between controls and individuals with TBI (Taylor et al., 2020). The multivariate D2 measure allowed for the discrimination of even mild TBI cases from controls and correlated significantly with cognitive scores. Similarly, using D2 combining both spatial (i.e., WM regions) and feature (i.e., different DTI metrics) dimensions led to improved detection between autistic and typically developing individuals compared to univariate approaches or to D2 computed by combining brain regions only (Dean et al., 2017). Associations between D2 and autism symptom severity were also reported in this study, providing additional evidence that D2 can serve as a behaviorally relevant measure of WM abnormality.

Other interesting implementations have used D2 to detect and characterize lesions. Gyebnár et al. (2019) combined DTI eigenvalues into a voxel-wise D2 measure between epilepsy patients and controls to detect cortical malformations in patients. Voxels were identified as belonging to a lesion if their D2 value exceeded a critical value calculated using Wilks’ criterion (Wilks, 1963), a criterion used for multivariate statistical outlier detection. In another implementation, D2 was employed to characterize the heterogeneity within WM lesions by computing the multivariate distance (combining T1-w, T2-w and PD-w signal intensities) between voxels in WM hyperintensities and those in normal appearing WM (NAWM) (Lindemer et al., 2015). D2 in WM hyperintensities progressed at a quicker rate in individuals who converted from mild cognitive impairment to Alzheimer’s disease (AD) compared to those who did not convert. Interestingly, the rate of change of WM hyperintensities volume (i.e., lesion load), a metric more commonly used (Bilello et al., 2015; Schmidt et al., 2005), did not differentiate converters from non-converters cross-sectionally and longitudinally, suggesting that a characterization of WM lesion heterogeneity through a multivariate framework was more informative than the volume of WM lesions (Lindemer et al., 2015).

### Limitations

There are some limitations of D2 computation as presented in MVComp. First, D2 itself is a squared measure, thus the directionality of the difference is non-specific. As it is currently implemented, it is not possible to determine whether a given subject’s features are higher or lower than the average, although this information can be easily extracted by comparing the subject’s voxel values or ROI means to the mean of the group average on a per-metric basis. Future studies could potentially address this limitation indirectly by integrating with studies that model ground-truth biophysical properties to better interpret differences and/or splitting groups based on expected direction of change. Then, the directions of deviations from the average could be hypothesized a priori.

D2 is a sensitive multivariate distance measure that has since found applications in various fields, such as classification, cluster analysis, and outlier detection. Our implementation makes use of the sensitivity of D2 to detect multivariate deviations in WM microstructure. This high sensitivity also means the method can be affected by registration inaccuracies and partial voluming (PV). Therefore, special attention must be paid to ensure optimal alignment across subjects and modalities (e.g., using directional information from dMRI to align WM tracts). Strict tissue type masking (e.g., using a high threshold on probabilistic segmentation images) can also be used to limit the amount of PV. However, this may result in a large number of excluded voxels, especially for low resolution images. Alternatively, the PV effect can be quantified and accounted for (e.g., González Ballester et al., 2002; Gyebnár et al., 2019). The latter option would be preferable if the D2 framework was used to detect tumors and estimate their volume, for instance.

Another limitation of D2 as presented in MVComp is that its use is restricted to continuous variables. However, more recent formulations of D2 allow for nominal and ordinal variables to be incorporated in the model, in addition to continuous variables (Barhen & Daudin, 1995; de Leon & Carrière, 2005). Future developments of MVComp could thus allow generalization of D2 to include mixed data types (e.g. WM, sex, or other grouping variable).

## 5. Conclusion

We introduce a new open-source tool for the computation of the Mahalanobis distance (D2), the MVComp (MultiVariate Comparisons) toolbox. D2 is a multivariate distance measure relative to a reference that inherently accounts for covariance between features. MVComp can be used in a wide range of neuroimaging implementations, at both the group and subject levels. In line with the current shift towards precision medicine, MVComp can be used to obtain personalized assessments of brain structure and function, which is essential in the study of brain conditions with high heterogeneity.

## Data and Code Availability

The data is openly available from the Human Connectome Project (https://www.humanconnectome.org/study/hcp-young-adult/document/1200-subjects-data-release) and the code of the MVComp toolbox is available at https://github.com/neuralabc/mvcomp (Tremblay et al., 2024).

## Author Contributions

**Stefanie A Tremblay**: Writing - Original Draft, Conceptualization, Data Curation, Methodology, Software, Validation, Visualization

**Zaki Alasmar**: Methodology, Software, Formal analysis, Validation, Conceptualization, Data Curation, Writing - Original Draft, Visualization

**Amir Pirhadi**: Methodology, Software, Validation, Data Curation

**Felix Carbonell**: Methodology, Writing - Review & Editing

**Yasser Iturria-Medina**: Methodology, Writing - Review & Editing, Conceptualization

**Claudine J Gauthier**: Supervision, Conceptualization, Writing - Review & Editing

**Christopher J Steele**: Supervision, Conceptualization, Methodology, Writing - Review & Editing, Software, Funding acquisition

## Supplementary Material

**Table 1.** Comparisons between subject(s) and a reference – Combining MRI metrics.

| MVComp function name | Description |
| --- | --- |
| `compute_average` | To compute the group average maps of each metric (will serve as the reference). |
| `feature_gen` | Apply to the reference group average maps to extract the feature matrix (`m_f_mat` of shape $n$ voxels × $n$ features), a mask vector (`mat_mask` of shape $n$ voxels) and a nibabel object of the mask (`mask_img`). |
| `norm_covar_inv` | To compute the covariance matrix (`s`) and its pseudoinverse (`pinv_s`) from the reference feature and mask matrices (`m_f_mat` and `mat_mask`). |
| `correlation_fig` | To generate a correlation matrix figure from the covariance matrix (`s`). For visualization. |
| `model_comp` | To calculate voxel-wise D2 between each subject contained in the provided `subject_ids` list and the reference (group average). Yields a D2 matrix of size number of voxels × number of subjects.<br>*For leave-one-out approach, set the `exclude_comp_from_mean_cov` option to True (the previous steps can be skipped in this case since a new covariance matrix is computed for each subject, within the `model_comp` function). |
| `dist_plot` | To produce D2 maps for every subject from the D2 matrix generated by `model_comp`. |
| `model_comp` with `return_raw` set to True | To extract features contribution to D2 in a region of interest. When `return_raw` is set to True, the function returns a 3D array of size (number of voxels) x (number of metrics) x (number of subjects). This information can then be summarized to obtain the % contribution of each metric for a group of subjects. |

**Table 2.** Comparisons between subject(s) and a reference – Combining spatial dimensions.

| MVComp function name | Description |
| --- | --- |
| `spatial_mvcomp` | To compute a D2 score between each subject and the reference from a matrix containing the data (e.g., mean FA in each WM tract) of all subjects ($n$ subjects x $n$ tracts). Returns a vector with a single D2 value per subject.<br>*For leave-one-out approach, set the `exclude_comp_from_mean_cov` option to True. |
| `spatial_mvcomp` with `return_raw` set to True | To extract features contribution to D2. If set to True, a 2D array of size (number of subjects) x (number of tracts) is returned. This information can then be summarized to obtain the relative importance of each tract to D2. |

**Table 3.** Comparisons within a single subject – Voxel-wise D2 resolution.

| MVComp function name | Description |
| --- | --- |
| `feature_gen` | Provide the path of the images (i.e., one image per metric) and the reference ROI mask to this function to extract the feature matrix (`m_f_mat` of shape $n$ voxels × $n$ features), a mask vector (`mat_mask` of shape $n$ voxels) and a nibabel object of the mask (`mask_img`). This function can also be used to extract the data inside the ROI of voxels to be evaluated. |
| `norm_covar_inv` | To compute the covariance matrix (`s`) and its pseudoinverse (`pinv_s`) from the reference feature and mask matrices (`m_f_mat` and `mat_mask`) . |
| `correlation_fig` | To generate a correlation matrix figure from the covariance matrix (`s`). For visualization. |
| `mah_dist_mat_2_roi` | To compute voxel-wise D2 between all voxels within a mask and a specific reference ROI. The user will need to provide a vector of data for the reference ROI (i.e., mean across voxels in the ROI for each metric), along with the feature matrix containing the data for the voxels to be evaluated. |
| `mah_dist_mat_2_roi` with `return_raw` set to True | To extract features' contributions. The output will be of shape (number of voxels) × (number of metrics). |

**Table 4.** Comparisons within a single subject – Voxel-voxel matrix D2 resolution.

| MVComp function name | Description |
| --- | --- |
| `voxel2voxel_dist` | To compute D2 between each voxel and all other voxels in a mask. Yields a symmetric 2-D matrix of size $n$ voxels × $n$ voxels containing D2 values between each pair of voxels. |

## References

1. Aboitiz, F., Scheibel, A. B., Fisher, R. S., & Zaidel, E. (1992). Fiber composition of the human corpus callosum. Brain Research, 598(1), 143–153. 10.1016/0006-8993(92)90178-C

2. Alexander-Bloch, A., Giedd, J. N., & Bullmore, E. (2013). Imaging structural co-variance between human brain regions. Nature Reviews Neuroscience, 14(5), Article 5. 10.1038/nrn3465

3. Alexopoulos, E. C. (2010). Introduction to Multivariate Regression Analysis. Hippokratia, 14(Suppl 1), 23–28.

4. Avants, B., Duda, J. T., Kim, J., Zhang, H., Pluta, J., Gee, J. C., & Whyte, J. (2008). Multivariate Analysis of Structural and Diffusion Imaging in Traumatic Brain Injury. Academic Radiology, 15(11), 1360–1375. 10.1016/j.acra.2008.07.007

5. Azzarito, M., Emmenegger, T. M., Ziegler, G., Huber, E., Grabher, P., Callaghan, M. F., Thompson, A., Friston, K., Weiskopf, N., Killeen, T., & Freund, P. (2023). Coherent, time-shifted patterns of microstructural plasticity during motor-skill learning. NeuroImage, 274, 120128. 10.1016/j.neuroimage.2023.120128

6. Barhen, A., & Daudin, J. J. (1995). Generalization of the Mahalanobis Distance in the Mixed Case. Journal of Multivariate Analysis, 53(2), 332–342. 10.1006/jmva.1995.1040

7. Basser, P. J., Mattiello, J., & LeBihan, D. (1994a). Estimation of the effective self-diffusion tensor from the NMR spin echo. Journal of Magnetic Resonance, Series B, 103(3), 247– 254.

8. Basser, P. J., Mattiello, J., & LeBihan, D. (1994b). MR diffusion tensor spectroscopy and imaging. Biophysical Journal, 66(1), 259–267. 10.1016/S0006-3495(94)80775-1

9. Bilello, M., Doshi, J., Nabavizadeh, S. A., Toledo, J. B., Erus, G., Xie, S. X., Trojanowski, J. Q., Han, X., & Davatzikos, C. (2015). Correlating Cognitive Decline with White Matter Lesion and Brain Atrophy Magnetic Resonance Imaging Measurements in Alzheimer’s Disease. Journal of Alzheimer’s Disease, 48(4), 987–994. 10.3233/JAD-150400

10. Calamante, F., Smith, R. E., Tournier, J.-D., Raffelt, D., & Connelly, A. (2015). Quantification of voxel-wise total fibre density: Investigating the problems associated with track-count mapping. NeuroImage, 117, 284–293. 10.1016/j.neuroimage.2015.05.070

11. Calhoun, V. d., Adali, T., Pearlson, G. d., & Pekar, J. j. (2001). A method for making group inferences from functional MRI data using independent component analysis. Human Brain Mapping, 14(3), 140–151. 10.1002/hbm.1048

12. Carbonell, F., Zijdenbos, A. P., & Bedell, B. J. (2020). Spatially Distributed Amyloid-β Reduces Glucose Metabolism in Mild Cognitive Impairment. Journal of Alzheimer’s Disease, 73(2), 543–557. 10.3233/JAD-190560

13. Carter, F., Anwander, A., Goucha, T., Adamson, H., Friederici, A. D., Lutti, A., Gauthier, C. J., Weiskopf, N., Bazin, P.-L., & Steele, C. J. (2022). Assessing Quantitative MRI Techniques using Multimodal Comparisons (p. 2022.02.10.479780). bioRxiv. 10.1101/2022.02.10.479780

14. Chamberland, M., Genc, S., Tax, C. M. W., Shastin, D., Koller, K., Raven, E. P., Cunningham, A., Doherty, J., van den Bree, M. B. M., Parker, G. D., Hamandi, K., Gray, W. P., & Jones, D. K. (2021). Detecting microstructural deviations in individuals with deep diffusion MRI tractometry. Nature Computational Science, 1(9), Article 9. 10.1038/s43588-021-00126-8

15. Chao, Y., Cho, K., Yeh, C., Chou, K., Chen, J., & Lin, C. (2009). Probabilistic topography of human corpus callosum using cytoarchitectural parcellation and high angular resolution diffusion imaging tractography. Human Brain Mapping, 30(10), 3172–3187. 10.1002/hbm.20739

16. Chen, C., Cao, X., & Tian, L. (2019). Partial least squares regression performs well in MRI-based individualized estimations. Frontiers in Neuroscience, 13, 1282.

17. Daducci, A., Canales-Rodríguez, E. J., Zhang, H., Dyrby, T. B., Alexander, D. C., & Thiran, J.-P. (2015). Accelerated Microstructure Imaging via Convex Optimization (AMICO) from diffusion MRI data. NeuroImage, 105, 32–44. 10.1016/j.neuroimage.2014.10.026

18. de Leon, A. R., & Carrière, K. C. (2005). A generalized Mahalanobis distance for mixed data. Journal of Multivariate Analysis, 92(1), 174–185. 10.1016/j.jmva.2003.08.006

19. Dean, D. C., Lange, N., Travers, B. G., Prigge, M. B., Matsunami, N., Kellett, K. A., Freeman, A., Kane, K. L., Adluru, N., Tromp, D. P. M., Destiche, D. J., Samsin, D., Zielinski, B. A., Fletcher, P. T., Anderson, J. S., Froehlich, A. L., Leppert, M. F., Bigler, E. D., Lainhart, J. E., & Alexander, A. L. (2017). Multivariate characterization of white matter heterogeneity in autism spectrum disorder. NeuroImage: Clinical, 14, 54–66. 10.1016/j.nicl.2017.01.002

20. Dhollander, T., Tabbara, R., Rosnarho-Tornstrand, J., Tournier, J.-D., Raffelt, D., & Connelly, A. (2021). Multi-tissue log-domain intensity and inhomogeneity normalisation for quantitative apparent fibre density. In Proc. ISMRM.

21. Ghorbani, H. (2019). MAHALANOBIS DISTANCE AND ITS APPLICATION FOR DETECTING MULTIVARIATE OUTLIERS. *Facta Universitatis*, Series: Mathematics and Informatics, 0, Article 0. 10.22190/FUMI1903583G

22. Glasser, M. F., & Essen, D. C. V. (2011). Mapping Human Cortical Areas In Vivo Based on Myelin Content as Revealed by T1-and T2-Weighted MRI. Journal of Neuroscience, 31(32), 11597–11616. 10.1523/JNEUROSCI.2180-11.2011

23. Glasser, M. F., Sotiropoulos, S. N., Wilson, J. A., Coalson, T. S., Fischl, B., Andersson, J. L., Xu, J., Jbabdi, S., Webster, M., Polimeni, J. R., Van Essen, D. C., & Jenkinson, M. (2013). The minimal preprocessing pipelines for the Human Connectome Project. NeuroImage, 80, 105–124. 10.1016/j.neuroimage.2013.04.127

24. González Ballester, M. Á., Zisserman, A. P., & Brady, M. (2002). Estimation of the partial volume effect in MRI. Medical Image Analysis, 6(4), 389–405. 10.1016/S1361-8415(02)00061-0

25. Guberman, G. I., Stojanovski, S., Nishat, E., Ptito, A., Bzdok, D., Wheeler, A. L., & Descoteaux, M. (2022). Multi-tract multi-symptom relationships in pediatric concussion. eLife, 11, e70450. 10.7554/eLife.70450

26. Guerrero-Gonzalez, J. M., Yeske, B., Kirk, G. R., Bell, M. J., Ferrazzano, P. A., & Alexander, A. L. (2022). Mahalanobis distance tractometry (MaD-Tract) – a framework for personalized white matter anomaly detection applied to TBI. NeuroImage, 260, 119475. 10.1016/j.neuroimage.2022.119475

27. Gyebnár, G., Klimaj, Z., Entz, L., Fabó, D., Rudas, G., Barsi, P., & Kozák, L. R. (2019). Personalized microstructural evaluation using a Mahalanobis-distance based outlier detection strategy on epilepsy patients’ DTI data – Theory, simulations and example cases. PLOS ONE, 14(9), e0222720. 10.1371/journal.pone.0222720

28. Hayasaka, S., Du, A.-T., Duarte, A., Kornak, J., Jahng, G.-H., Weiner, M. W., & Schuff, N. (2006). A non-parametric approach for co-analysis of multi-modal brain imaging data: Application to Alzheimer’s disease. NeuroImage, 30(3), 768–779. 10.1016/j.neuroimage.2005.10.052

29. Hofer, S., & Frahm, J. (2006). Topography of the human corpus callosum revisited— Comprehensive fiber tractography using diffusion tensor magnetic resonance imaging. NeuroImage, 32(3), 989–994. 10.1016/j.neuroimage.2006.05.044

30. Hotelling, H. (1947). Multivariate quality control. Techniques of Statistical Analysis.

31. Iturria-Medina, Y., Carbonell, F. M., Sotero, R. C., Chouinard-Decorte, F., & Evans, A. C. (2017). Multifactorial causal model of brain (dis)organization and therapeutic intervention: Application to Alzheimer’s disease. NeuroImage, 152, 60–77. 10.1016/j.neuroimage.2017.02.058

32. Jeurissen, B., Tournier, J.-D., Dhollander, T., Connelly, A., & Sijbers, J. (2014). Multi-tissue constrained spherical deconvolution for improved analysis of multi-shell diffusion MRI data. NeuroImage, 103, 411–426. 10.1016/j.neuroimage.2014.07.061

33. Khedher, L., Ramírez, J., Górriz, J. M., Brahim, A., & Segovia, F. (2015). Early diagnosis of Alzheimer’s disease based on partial least squares, principal component analysis and support vector machine using segmented MRI images. Neurocomputing, 151, 139–150. 10.1016/j.neucom.2014.09.072

34. Kritzman, M., & Li, Y. (2010). Skulls, Financial Turbulence, and Risk Management. Financial Analysts Journal, 66(5), 30–41. 10.2469/faj.v66.n5.3

35. Kulikova, S., Hertz-Pannier, L., Dehaene-Lambertz, G., Buzmakov, A., Poupon, C., & Dubois, J. (2015). Multi-parametric evaluation of the white matter maturation. Brain Structure and Function, 220(6), 3657–3672. 10.1007/s00429-014-0881-y

36. Lindemer, E. R., Salat, D. H., Smith, E. E., Nguyen, K., Fischl, B., & Greve, D. N. (2015). White matter signal abnormality quality differentiates mild cognitive impairment that converts to Alzheimer’s disease from nonconverters. Neurobiology of Aging, 36(9), 2447–2457. 10.1016/j.neurobiolaging.2015.05.011

37. Mahalanobis, P. C. (1936). On the generalized distance in statistics.

38. Marquand, A. F., Rezek, I., Buitelaar, J., & Beckmann, C. F. (2016). Understanding Heterogeneity in Clinical Cohorts Using Normative Models: Beyond Case-Control Studies. Biological Psychiatry, 80(7), 552–561. 10.1016/j.biopsych.2015.12.023

39. Naylor, M. G., Cardenas, V. A., Tosun, D., Schuff, N., Weiner, M., & Schwartzman, A. (2014). Voxelwise multivariate analysis of multimodality magnetic resonance imaging. Human Brain Mapping, 35(3), 831–846. 10.1002/hbm.22217

40. Nestor, P. G., O’Donnell, B. F., McCarley, R. W., Niznikiewicz, M., Barnard, J., Jen Shen, Z., Bookstein, F. L., & Shenton, M. E. (2002). A new statistical method for testing hypotheses of neuropsychological/MRI relationships in schizophrenia: Partial least squares analysis. Schizophrenia Research, 53(1), 57–66. 10.1016/S0920-9964(00)00171-7

41. Novikov, D. S., Kiselev, V. G., & Jespersen, S. N. (2018). On modeling. Magnetic Resonance in Medicine, 79(6), 3172–3193. 10.1002/mrm.27101

42. Owen, T. W., de Tisi, J., Vos, S. B., Winston, G. P., Duncan, J. S., Wang, Y., & Taylor, P. N. (2021). Multivariate white matter alterations are associated with epilepsy duration. The European Journal of Neuroscience, 53(8), 2788–2803. 10.1111/ejn.15055

43. Patenaude, B., Smith, S. M., Kennedy, D. N., & Jenkinson, M. (2011). A Bayesian model of shape and appearance for subcortical brain segmentation. NeuroImage, 56(3), 907–922. 10.1016/j.neuroimage.2011.02.046

44. Plitman, E., Ochi, R., Patel, R., Tsugawa, S., Tarumi, R., Honda, S., Matsushita, K., Fujii, S., Uchida, H., & Mimura, M. (2020). Using Non-Negative Matrix Factorization to Examine Treatment Resistance and Response in Patients With Schizophrenia: A Multimodal Imaging Study. Biological Psychiatry, 87(9), S350.

45. Raffelt, D., Dhollander, T., Tournier, J.-D., Tabbara, R., Smith, R. E., Pierre, E., & Connelly, A. (2017). Bias Field Correction and Intensity Normalisation for Quantitative Analysis of Apparent Fibre Density. In Proc. ISMRM.

46. Raffelt, D., Tournier, J.-D., Fripp, J., Crozier, S., Connelly, A., & Salvado, O. (2011). Symmetric diffeomorphic registration of fibre orientation distributions. NeuroImage, 56(3), 1171– 1180. 10.1016/j.neuroimage.2011.02.014

47. Raffelt, D., Tournier, J.-D., Rose, S., Ridgway, G. R., Henderson, R., Crozier, S., Salvado, O., & Connelly, A. (2012). Apparent Fibre Density: A novel measure for the analysis of diffusion-weighted magnetic resonance images. NeuroImage, 59(4), 3976–3994. 10.1016/j.neuroimage.2011.10.045

48. Raffelt, D., Tournier, J.-D., Smith, R. E., Vaughan, D. N., Jackson, G., Ridgway, G. R., & Connelly, A. (2017). Investigating white matter fibre density and morphology using fixel-based analysis. NeuroImage, 144, 58–73. 10.1016/j.neuroimage.2016.09.029

49. Schmidt, R., Ropele, S., Enzinger, C., Petrovic, K., Smith, S., Schmidt, H., Matthews, P. M., & Fazekas, F. (2005). White matter lesion progression, brain atrophy, and cognitive decline: The Austrian stroke prevention study. Annals of Neurology, 58(4), 610–616. 10.1002/ana.20630

50. Seidlitz, J., Váša, F., Shinn, M., Romero-Garcia, R., Whitaker, K. J., Vértes, P. E., Wagstyl, K., Kirkpatrick Reardon, P., Clasen, L., Liu, S., Messinger, A., Leopold, D. A., Fonagy, P., Dolan, R. J., Jones, P. B., Goodyer, I. M., NSPN Consortium, Raznahan, A., & Bullmore, E. T. (2018). Morphometric Similarity Networks Detect Microscale Cortical Organization and Predict Inter-Individual Cognitive Variation. Neuron, 97(1), 231–247.e7. 10.1016/j.neuron.2017.11.039

51. Smith, R. E., Tournier, J.-D., Calamante, F., & Connelly, A. (2012). Anatomically-constrained tractography: Improved diffusion MRI streamlines tractography through effective use of anatomical information. NeuroImage, 62(3), 1924–1938. 10.1016/j.neuroimage.2012.06.005

52. Smith, R. E., Tournier, J.-D., Calamante, F., & Connelly, A. (2013). SIFT: Spherical-deconvolution informed filtering of tractograms. NeuroImage, 67, 298–312. 10.1016/j.neuroimage.2012.11.049

53. Smith, S. M. (2002). Fast robust automated brain extraction. Human Brain Mapping, 17(3), 143–155. 10.1002/hbm.10062

54. Smith, S. M., Jenkinson, M., Woolrich, M. W., Beckmann, C. F., Behrens, T. E., Johansen-Berg, H., Bannister, P. R., De Luca, M., Drobnjak, I., & Flitney, D. E. (2004). Advances in functional and structural MR image analysis and implementation as FSL. Neuroimage, 23, S208–S219.

55. Tardif, C. L., Gauthier, C. J., Steele, C. J., Bazin, P.-L., Schäfer, A., Schaefer, A., Turner, R., & Villringer, A. (2016). Advanced MRI techniques to improve our understanding of experience-induced neuroplasticity. Neuroimage, 131, 55–72.

56. Taubert, M., Villringer, A., & Ragert, P. (2012). Learning-Related Gray and White Matter Changes in Humans: An Update. The Neuroscientist, 18(4), 320–325. 10.1177/1073858411419048

57. Taylor, P. N., Silva, N. M. da, Blamire, A., Wang, Y., & Forsyth, R. (2020). Early deviation from normal structural connectivity: A novel intrinsic severity score for mild TBI. Neurology, 94(10), e1021–e1026. 10.1212/WNL.0000000000008902

58. Thiebaut de Schotten, M., & Forkel, S. J. (2022). The emergent properties of the connected brain. Science, 378(6619), 505–510. 10.1126/science.abq2591

59. Tournier, J.-D., Smith, R., Raffelt, D., Tabbara, R., Dhollander, T., Pietsch, M., Christiaens, D., Jeurissen, B., Yeh, C.-H., & Connelly, A. (2019). MRtrix3: A fast, flexible and open software framework for medical image processing and visualisation. NeuroImage, 116137.

60. Tremblay, S.A., Alasmar, Z., Pirhadi, A., Carbonell, F., Iturria-Medina, Y., Gauthier, C., Steele, C.J. (2024) mvcomp: Multivariate Comparisons using Whole-brain and ROI-derived Metrics from MRI, v0.9.2, Zenodo, doi:10.5281/zenodo.10713027

61. Tustison, N. J., Avants, B. B., Cook, P. A., Yuanjie Zheng, Egan, A., Yushkevich, P. A., & Gee, J. C. (2010). N4ITK: Improved N3 Bias Correction. IEEE Transactions on Medical Imaging, 29(6), 1310–1320. 10.1109/TMI.2010.2046908

62. Uddin, M. N., Figley, T. D., Solar, K. G., Shatil, A. S., & Figley, C. R. (2019). Comparisons between multi-component myelin water fraction, T1w/T2w ratio, and diffusion tensor imaging measures in healthy human brain structures. Scientific Reports, 9(1), Article 1. 10.1038/s41598-019-39199-x

63. Van Essen, D. C., Smith, S. M., Barch, D. M., Behrens, T. E. J., Yacoub, E., Ugurbil, K., & WU-Minn HCP Consortium. (2013). The WU-Minn Human Connectome Project: An overview. NeuroImage, 80, 62–79. 10.1016/j.neuroimage.2013.05.041

64. Vandekar, S. N., Shinohara, R. T., Raznahan, A., Hopson, R. D., Roalf, D. R., Ruparel, K., Gur, R. C., Gur, R. E., & Satterthwaite, T. D. (2016). Subject-level measurement of local cortical coupling. NeuroImage, 133, 88–97. 10.1016/j.neuroimage.2016.03.002

65. Veraart, J., Sijbers, J., Sunaert, S., Leemans, A., & Jeurissen, B. (2013). Weighted linear least squares estimation of diffusion MRI parameters: Strengths, limitations, and pitfalls. NeuroImage, 81, 335–346. 10.1016/j.neuroimage.2013.05.028

66. Whitaker, K. J., Vértes, P. E., Romero-Garcia, R., Váša, F., Moutoussis, M., Prabhu, G., Weiskopf, N., Callaghan, M. F., Wagstyl, K., Rittman, T., Tait, R., Ooi, C., Suckling, J., Inkster, B., Fonagy, P., Dolan, R. J., Jones, P. B., Goodyer, I. M., the NSPN Consortium, & Bullmore, E. T. (2016). Adolescence is associated with genomically patterned consolidation of the hubs of the human brain connectome. Proceedings of the National Academy of Sciences, 113(32), 9105–9110. 10.1073/pnas.1601745113

67. Wilks, S. S. (1963). Multivariate Statistical Outliers. Sankhyā: The Indian Journal of Statistics, Series A (1961-2002), 25(4), 407–426.

68. Wolfers, T., Doan, N. T., Kaufmann, T., Alnæs, D., Moberget, T., Agartz, I., Buitelaar, J. K., Ueland, T., Melle, I., Franke, B., Andreassen, O. A., Beckmann, C. F., Westlye, L. T., & Marquand, A. F. (2018). Mapping the Heterogeneous Phenotype of Schizophrenia and Bipolar Disorder Using Normative Models. JAMA Psychiatry, 75(11), 1146–1155. 10.1001/jamapsychiatry.2018.2467

69. Xiang, S., Nie, F., & Zhang, C. (2008). Learning a Mahalanobis distance metric for data clustering and classification. Pattern Recognition, 41(12), 3600–3612. 10.1016/j.patcog.2008.05.018

70. Yang, W., Lui, R. L. M., Gao, J.-H., Chan, T. F., Yau, S.-T., Sperling, R. A., & Huang, X. (2011). Independent Component Analysis-Based Classification of Alzheimer’s Disease MRI Data. Journal of Alzheimer’s Disease, 24(4), 775–783. 10.3233/JAD-2011-101371

71. Young, K., Govind, V., Sharma, K., Studholme, C., Maudsley, A. A., & Schuff, N. (2010). Multivariate statistical mapping of spectroscopic imaging data. Magnetic Resonance in Medicine, 63(1), 20–24. 10.1002/mrm.22190

72. Zhang, H., Schneider, T., Wheeler-Kingshott, C. A., & Alexander, D. C. (2012). NODDI: Practical in vivo neurite orientation dispersion and density imaging of the human brain. NeuroImage, 61(4), 1000–1016. 10.1016/j.neuroimage.2012.03.072

73. Zhang, Y., Brady, M., & Smith, S. (2001). Segmentation of brain MR images through a hidden Markov random field model and the expectation-maximization algorithm. IEEE Transactions on Medical Imaging, 20(1), 45–57. 10.1109/42.906424

